# Perinatal Morphine Exposure Induces Long-Term Changes in the Intestinal Microbiota of Male and Female Rats

**DOI:** 10.1101/2023.09.20.558694

**Authors:** Hannah J. Harder, Charlène J.G. Dauriat, Benoit Chassaing, Anne Z. Murphy

## Abstract

The increased use of opioids by women of reproductive age has resulted in a dramatic rise in number of infants exposed to opioids *in utero*. Although perinatal opioid exposure (POE) has been associated with an elevated risk of infection and hospitalization later in life, the mechanism(s) by which opioids influence immune development and maturation is not fully elucidated. Alterations in the intestinal microbiota composition, which leads to changes in immune training and maturation, could be at play. Chronic opioid use in adults is associated with a proinflammatory and pathogenic microbiota composition; therefore, we hypothesized here that *in utero* morphine exposure could negatively affect intestinal microbiota composition, leading to alterations in immune system function. We report that a clinically-relevant model of perinatal opioid exposure, in rats, induces profound intestinal microbiota dysbiosis that is maintained into adulthood. Furthermore, microbial maturity was reduced in morphine-exposed offspring. This suggests that increased risk of infection observed in children exposed to opioids during gestation may be a consequence of microbiota alterations with downstream impact on immune system development. Further investigation of how perinatal morphine induces dysbiosis will be critical to the development of early life interventions designed to ameliorate the increased risk of infection observed in these children.

## Introduction

Maternal opioid use has risen by 131% since 2010 (Hirai et al., 2021). As a result, a staggering 6.3 per 1000 infants were born in the United States in 2020 experiencing neonatal opioid withdrawal syndrome (NOWS) (Healthcare Cost and Utilization Project, 2022; Hirai et al., 2021). Symptoms associated with NOWS mirror those typically observed in adults and are predominated by gastrointestinal problems, including constipation, nausea, abdominal pain, and diarrhea (Kocherlakota, 2014; Leppert, 2015). Opioid use and misuse have also been linked to gut microbiota dysbiosis, or an altered composition of the commensal bacteria found in the gastrointestinal tract (Akbarali and Dewey, 2019). Both opioid-induced gastrointestinal issues and dysbiosis are a result of opioid action on µ-opioid receptors in the gut, which decrease motility and allow for bacterial overgrowth (Akbarali and Dewey, 2019), predisposing organisms to inflammation, infection, and disease (Round and Mazmanian, 2009; Valdes et al., 2018). Although the microbiota is an essential immune stimulator in early life, very few, if any, clinical studies have investigated gut microbiota composition in infants born with NOWS. It is theorized that exposure to opioids *in utero* leads to a dysbiotic gut microbiota, which is maintained across the lifespan, influencing immune system development (Maguire and Gröer, 2016). Importantly, children exposed to opioids *in utero* have an increased risk of infection (Arter et al., 2021; Uebel et al., 2015; Witt et al., 2017), consistent with impaired immune system development.

Chronic opioid exposure in adulthood generally promotes a proinflammatory gut microbiota composition as a result of decreased intestinal transit and increased gut permeability. Although individual studies report different effects on microbiota alpha or beta diversity, it is clear that systemic opioid exposure in adulthood alters gut microbiota composition (Akbarali and Dewey, 2019). Alpha diversity includes richness, referring to the number of overall microbial species present in a sample, as well as evenness, referring to the distribution of abundance of all species in the sample. Previous studies on the influence of opioids on alpha diversity report conflicting results (Lee et al., 2018; Ren and Lotfipour, 2022; Zhang et al., 2021), likely due to differences in the type of opioid administered, timing and length of dosing schedule, as well as timepoints sampled after opioid exposure; however, the majority report a decrease in alpha diversity with chronic opioid use (Cruz-Lebrón et al., 2021; Gicquelais et al., 2020; Ren and Lotfipour, 2022; Sharma et al., 2020; Wang et al., 2018). Reduced alpha diversity is associated with a variety of negative health outcomes and neurological diseases (Cryan et al., 2020), including autism spectrum disorder, Alzheimer’s disease, and epilepsy. Although no clinical studies have investigated alpha diversity in children exposed to opioids *in utero*, this suggests that alpha diversity may be altered long-term, potentially predisposing these children to future health challenges.

Beta diversity, or the differences in the composition of the gut microbiota between groups, is also altered by chronic opioid exposure in adults. Commonly reported changes in intestinal microbiota composition include expansion of proinflammatory genera of bacteria, including *Staphylococcus, Sutterella, Enterococcus*, and loss of probiotic or protective microbes, including *Lactobacillus*, Lachnospiraceae, and Ruminococcaceae (Fürst et al., 2020). Opioid exposure also reduces the abundance of bile-deconjugating microbes, leading to lower levels of anti-inflammatory bile acids (Banerjee et al., 2016; Wang et al., 2018). Decreases in bile acid levels are associated with gut barrier disruption and inflammation, further altering intestinal microbiota composition through decreased secretion of antimicrobial peptides.

The above-mentioned studies focused primarily on chronic opioid use in adults. Thus, there is limited preclinical evidence on the impact of POE on the gut microbiota composition. These studies have associated POE with altered alpha (Grecco et al., 2021) and beta diversity (Abu et al., 2021; Grecco et al., 2021; Lyu et al., 2022). Importantly, many of the bacterial taxa identified as differentially abundant after POE are associated with bile acid production (including Ruminococcaceae, Rikenellaceae, Erysipelotrichaceae, Lachnospiraceae, and Allobaculum), suggesting a possible mechanism by which *in utero* opioid exposure influences gut inflammation and barrier function.

Exposure to opioids is associated with a pathogenic profile of the gut microbiota, promoting immune activation, inflammation, and cytokine release. Previous work has associated chronic opioid exposure with increased morbidity and mortality from infection, which was directly related to gut microbiota dysbiosis (Babrowski et al., 2012; Meng et al., 2015; Mora et al., 2011; Wang et al., 2020). These findings are consistent with our previous study in which the febrile and neuroinflammatory response induced by lipopolysaccharide administration is potentiated in perinatally-exposed male and female rats (Harder et al., 2023). However, the mechanism by which POE alters immune function is unknown. Thus, the present studies were conducted to test the hypothesis that rats perinatally exposed to morphine would have long-term alterations in gut microbiota composition that may contribute to the potentiated response to an immune challenge.

## Methods

### Experimental subjects

Female Sprague Dawley rats (approximately two months of age; Charles River Laboratories, Boston, MA) were used to generate offspring. Same-sex pairs or groups of three were co-housed in Optirat GenII individually ventilated cages (Animal Care Systems, Centennial, Colorado, USA) with corncob bedding on a 12:12 hours light/dark cycle (lights on at 8:00 AM). Food (Lab Diet 5001 or Lab Diet 5015 for breeding pairs, St. Louis, MO, USA) and water were provided *ad libitum* throughout the experiment, except during testing. All studies were approved by the Institutional Animal Care and Use Committee at Georgia State University and performed in compliance with the National Institutes of Health Guide for the Care and Use of Laboratory Animals. All efforts were made to reduce the number of rats used in these studies and minimize pain and suffering.

### Perinatal opioid exposure paradigm

Briefly, female Sprague Dawley rats were implanted with iPrecio® SMP-200 microinfusion minipumps at postnatal day 60 (P60) under isoflurane anesthesia. Pumps were programmed to deliver 10-16 mg/kg three times a day. One week after morphine initiation, females were paired with sexually-experienced males for two weeks to induce pregnancy. Morphine exposure to the dams continued throughout gestation. Dams continued to receive morphine after parturition, such that pups received morphine through maternal milk. Beginning at P5, morphine dosage was decreased by 2 mg/kg daily until P7, when morphine administration was discontinued. A separate cohort of rats were implanted with pumps filled with sterile saline to control for the stress of surgery and pump refilling.

### Fecal sample collection

Fecal samples were collected from both dams and offspring. For dams, rats were isolated into clean containers for fresh fecal collection across the dosing paradigm (10 mg/kg, 12 mg/kg, 14 mg/kg, and 16 mg/kg morphine or the equivalent for vehicle dams). For offspring, morphine-exposed (MOR) and vehicle (VEH) male and female rats were isolated into clean containers for fresh fecal collection at P21, P28, P42, P56, and P70. Due to small sample sizes at P21 and P28, fecal samples were combined per cage at those timepoints (all cages were treatment- and sex-matched). Fecal samples were collected into sterile tubes and promptly frozen at −20°C until sequencing. See Figure 1 for a description of the dosing protocol and sampling timepoints.

**Figure 1.**
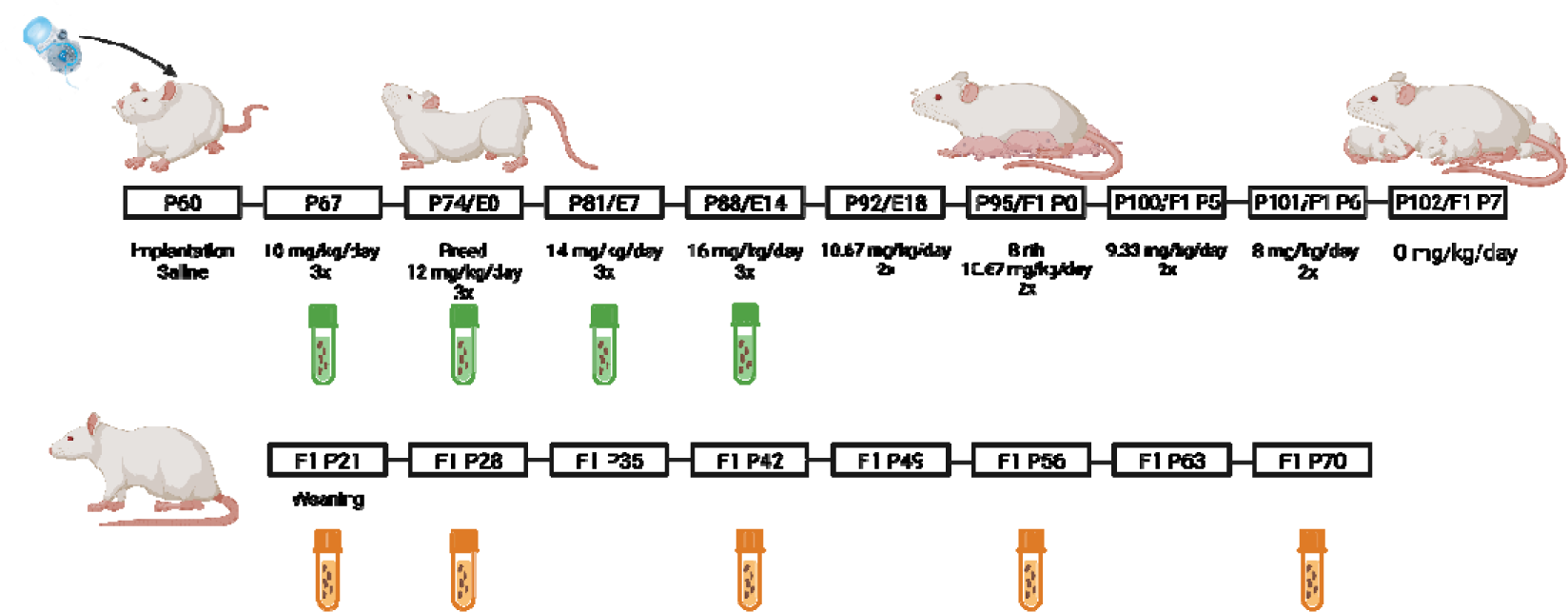
Timeline of opioid dosing and sample collection. Green tubes represent fecal sample collection from dams; orange tubes represent collection from offspring.

### Microbiota analysis by 16 S rRNA gene sequencing

16S rRNA gene amplification and sequencing were conducted using the Illumina MiSeq technology following the protocol of the Earth Microbiome Project (https://earthmicrobiome.org/protocols-and-standards/). Bulk DNA was extracted from frozen feces using a QIAamp 96 PowerFecal QIAcube HT kit (Qiagen Laboratories) with mechanical disruption (Qiagen TissueLyser II). The 16S rRNA genes, region V4, were PCR amplified from each sample using a composite forward primer and a reverse primer containing a unique 12-base barcode, designed using the Golay error-correcting scheme, which was used to tag PCR products from respective samples (Caporaso et al., 2012; Naimi et al., 2021). The present study used the forward primer 515F 5’-*AATGATACGGCGACCACCGAGATCTACACGCT*XXXXXXXXXXXX**TATGGTAATT*GT***<u>GTGYCAGCMGCCGCGGTAA</u>-3’:

1. the italicized sequence is the 5’ Illumina adapter;
2. the 12 X sequence is the Golay barcode;
3. the bold sequence is the primer pad;
4. the italicized and bold sequence is the primer linker;
5. and the underlined sequence is the conserved bacterial primer 515F.

The reverse primer 806R used was 5’-*CAAGCAGAAGACGGCATACGAGAT***AGTCAGCCAG*CC*** GGACTACNVGGGTWTCTAAT-3’:

1. the italicized sequence is the 3’ reverse complement sequence of the Illumina adapter;
2. the bold sequence is the primer pad;
3. the italicized and bold sequence is the primer linker;
4. and the underlined sequence is the conserved bacterial primer 806R.

PCR reactions consisted of Hot Master PCR mix (Quantabio, Beverly, MA, USA), 0.2 μM of each primer, and 10–100 ng template; reaction conditions were 3 min at 95°C, followed by 30 cycles of 45 s at 95°C, 60 s at 50*°*C, and 90 s at 72°C on a Biorad thermocycler. Products were then visualized by gel electrophoresis and quantified using Quant-iT PicoGreen dsDNA assay (Clariostar Fluorescence Spectrophotometer). A master DNA pool was generated in equimolar ratios, subsequently purified with Ampure magnetic purification beads (Agencourt, Brea, CA, USA) and sequenced using an Illumina MiSeq sequencer (paired-end reads, 2 × 250 bp) at the Genom’IC platform (INSERM U1016, Paris, France).

### 16S rRNA gene sequence analysis

16S rRNA sequences were analyzed using QIIME2—version 2019, as previously described (Bolyen et al., 2019; Naimi et al., 2021). Sequences were demultiplexed and quality filtered using the Dada2 method with QIIME2 default parameters in order to detect and correct Illumina amplicon sequence data, and a table of QIIME2 artifacts was generated. A tree was next generated, using the align-to-tree-mafft-fasttree command, for phylogenetic diversity analyses; alpha and beta diversity analyses were computed using the core-metrics-phylogenetic command. To compare species richness and evenness, three measures of alpha diversity were calculated: Pielou’s evenness, Shannon’s index, and Simpson’s index. Principal coordinate analysis (PCoA) plots were used to assess the variation between the experimental groups (beta diversity) as generated with the R package qiime2R v0.99.6, along with PERMANOVA for statistical analysis of treatment differences (as implemented in Qiime2 using the beta-group-significance command). Two measures of beta diversity were calculated: Bray-Curtis dissimilarity and unweighted Unifrac distance. For taxonomy analysis, amplicon sequence variants (ASVs) were assigned to operational taxonomic units (OTUs) with a 99% threshold of pairwise identity to the Greengenes reference database 13_8.

### Microbiota analysis using Maaslin2 to identify differentially abundant OTUs

To determine OTUs differentially expressed across metadata categories (i.e., drug treatment, sex, and/or age), we utilized the R package MaAsLin2 v1.10.0 (Mallick et al., 2021). MaAsLin2 utilizes general linear models, metadata, and rarefied taxonomic data to compare abundance of individual OTUs between metadata categories. Abundance was compared both for MOR vs. VEH dams across the dosing schedule (10 mg/kg, 12 mg/kg, 14 mg/kg, 16 mg/kg morphine or equivalent for saline). We also compared MOR vs. VEH offspring across age (P21, P28, P42, P56, and P70) separately by sex. For offspring, random effects of litter and experimental cohort were also included. For analysis, min_prevalence was set to 0.1 (the minimum percentage of samples an OTU must be detected at minimum abundance for further analysis; 10%) and min_abundance was set to 0.0001 (the minimum abundance for each OTU; non-zero). All other parameters used default values. Individual OTUs were compared between treatment, sex, and age to generate coefficients of the individual comparisons, along with p values to represent significance. Those p values were then corrected for multiple comparisons via the Benjamini-Hochberg false discovery rate correction to generate q values. Any differences with q<0.05 were considered significant. Effect sizes were calculated via -log(qvalue), multiplied by the sign of the coefficient of the difference between treatment groups. Positive effect sizes represent higher abundance in MOR rats. All comparisons were made to the sex- and age-matched vehicle controls.

### Microbiota analysis using maturity-index in Qiime2

As we examined gut microbiota composition in MOR and VEH rats across multiple ages (P21, P28, P42, P56, and P70), we also investigated “microbial maturity” using the maturity-index function in Qiime2. By comparing gut microbiota composition over time in a reference group (in this case, VEH rats), maturity-index generates a regression model to predict age as a function of composition and then calculates maturity index z-scores (MAZ scores) to compare relative microbial maturity across groups. Lower MAZ scores represent relatively “immature microbial ecosystem. We also utilized the MAZ scores in a linear mixed-effects model, as implemented in Qiime2 with the longitudinal linear-mixed-effects function, in order to determine whether treatment or sex was a significant predictor of maturity z-score.

### Experimental design and statistical analysis

Female rats were randomly assigned to the morphine or vehicle condition. All experiments included both male and female offspring to investigate potential sex-specific effects of perinatal opioid exposure on gut microbiota composition. No differences were observed between rats of different litters in the same drug exposure group (i.e., MOR and VEH); therefore, individual rats from multiple litters served as a single sample count (3 VEH litters, 3 MOR litters). Data was collected from two separate rounds of rats. All analyses were completed blinded to treatment group.

Alpha diversity was analyzed using mixed models with Greenhouse-Geisser correction as implemented in GraphPad Prism 9.1.0 (Motulsky, 2023) to identify significant effects of treatment and dose/age; p<0.05 was considered significant. Sidak’s post-hoc tests were conducted to determine significant mean differences between *a priori* specified groups.

## Results

### Maternal alpha and beta diversity

As maternal microbiota could impact the offspring’s microbiota establishment, we first investigated the direct effects of morphine exposure on female rats before pregnancy and across gestation. Previous studies suggest that alterations in maternal microbiota during gestation can lead to long-term changes in offspring’s gut microbiota composition and immunity (Nyangahu et al., 2018), suggesting that maternal microbiota dysbiosis has a potential influence on offspring’s future health. Three specific indexes of alpha diversity were analyzed at each morphine dose (10, 12, 14, and 16 mg/kg) and compared to time-matched vehicle samples: Pielou’s evenness (Figure 2A), Shannon’s index (Figure 2B), and Simpson’s index (Figure 2C). No significant effects of treatment nor dose were found in any of the three alpha diversity metrics (see Table 1). Although not significant, alpha diversity generally decreased across pregnancy in both groups, suggesting that the effects of pregnancy may overshadow any effects of morphine.

**Figure 2.**
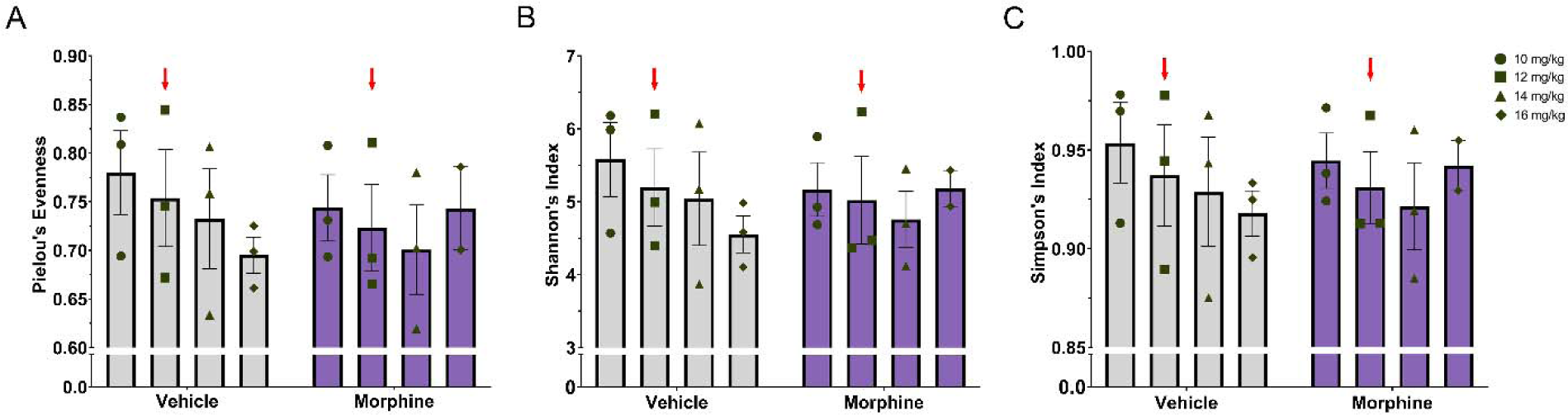
Alpha diversity does not differ between morphine- and vehicle-treated dams. **A.** No differences in Pielou’s evenness (**A**), Shannon’s index (**B**), or Simpson’s index (**C**). Red arrows indicate the initiation of breeding. N_VEH_ = 3, N_MOR_ = 3.

**Table 1.** F values for alpha diversity metrics for dams and offspring. * = significant at p<0.05.

|  | **Alpha Diversity Metric** | **Treatment** | **Dose/Age** | **Interaction** |
| --- | --- | --- | --- | --- |
| **Dam** | Pielou's Evenness | F(1,4)=0.05242,<br>p=0.8285 | F(2.350,8.618)=1.223,<br>p=0.3484 | F(3,11)=1.135,<br>p=0.3776 |
|  | Shannon's Index | F(1,4)=0.01303,<br>p=0.9146 | F(1.964,7.202)=0.8796,<br>p=0.4534 | F(3,11)=0.7716,<br>p=0.5337 |
|  | Simpson's Index | F(1,4)=0.0007879,<br>p=0.9790 | F(2.123,7.786)=1.546,<br>p=0.2733 | F(3,11)=0.8077,<br>p=0.5155 |
| **Offspring** | Pielou's Evenness | F(1,55)=0.2744,<br>p=0.6025 | F(3.158,72.61)=1.991,<br>p=0.1198 | F(4,92)=2.259,<br>p=0.0687 |
|  | Shannon's Index | F(1,55)=3.264,<br>p=0.0763 | F(3.086,70.97)=2.451,<br>p=0.0688 | F(4,92)=3.152,<br>p=0.0178* |
|  | Simpson's Index | F(1,55)=2.470,<br>p=0.1218 | F(3.387,77.90)=1.366,<br>p=0.2573 | F(4,92)=2.263,<br>p=0.0683 |

We next investigated beta diversity in order to compare intestinal microbiota composition between VEH and MOR dams. Two different measures of beta diversity were computed: Bray-Curtis dissimilarity and unweighted Unifrac distances. Bray-Curtis is a non-phylogenetic and weighted measure, while unweighted Unifrac is a phylogenetic and unweighted measure. PERMANOVA was used to determine if beta diversity significantly differed between VEH and MOR dams. Significant treatment effects were observed in both Bray-Curtis dissimilarity (p=0.019; Figure 3A) and unweighted Unifrac distances (p=0.042; Figure 3B), suggesting altered gut microbiota composition as a result of morphine treatment.

**Figure 3.**
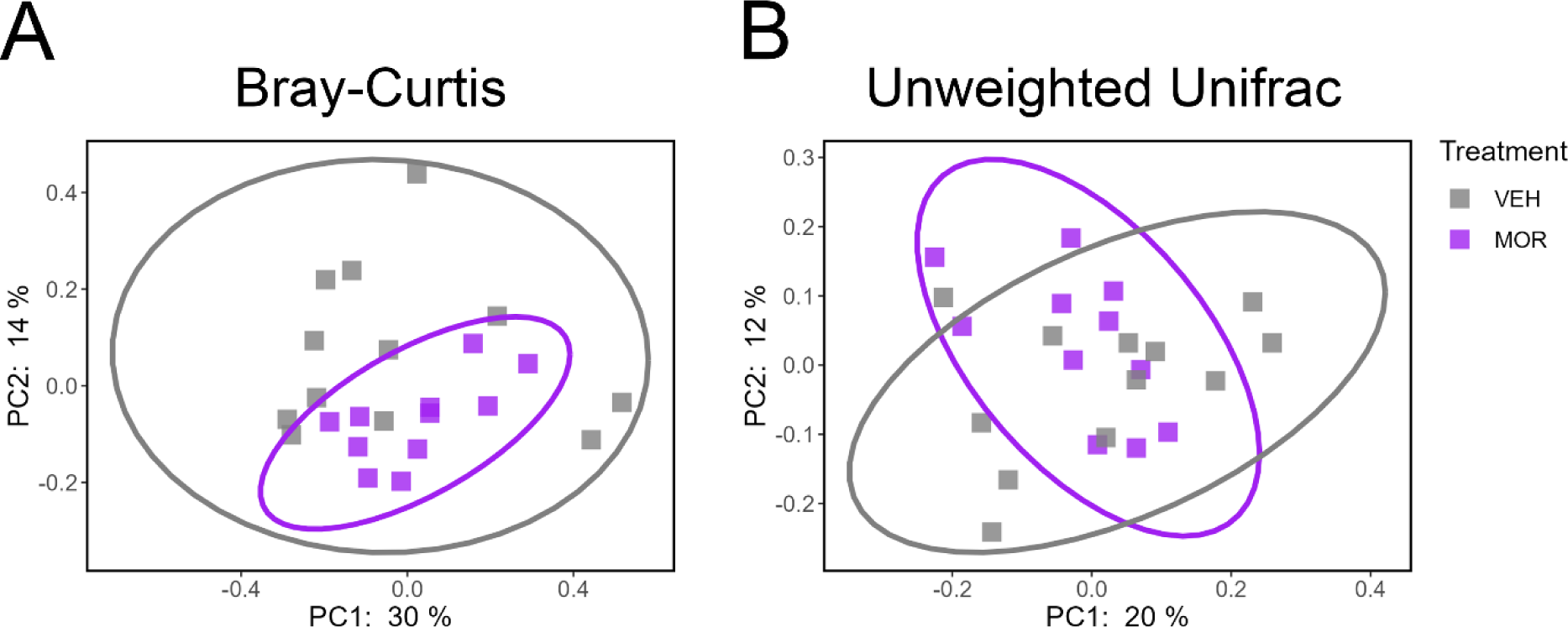
Morphine-exposed dams have significantly altered beta diversity in comparison to vehicle dams. **A.** Significant clustering on the Bray-Curtis dissimilarity measure based on treatment. **B.** Significant clustering of unweighted Unifrac distances based on treatment. N_VEH_ = 3, N_MOR_ = 3.

We next examined which OTUs were significantly impacted in their relative abundance by morphine exposure. For this purpose, we used MaAsLin2 to compare abundance between VEH and MOR dams at all four timepoints/morphine doses investigated. At 10 mg/kg, when morphine was first initiated and prior to breeding, two OTUs were significantly increased in MOR dams: an unidentified Ruminococcaceae member and the genus *Oscillospira*. Both the Ruminococcaceae family and its genus *Oscillospira* are generally considered probiotic and involved in both bile acid and short-chain fatty acid synthesis (Fürst et al., 2020; Yang et al., 2021). No other OTUs were significantly altered at any other time, suggesting that opioids may have larger effects at initiation vs. maintenance.

### Offspring alpha and beta diversity

We next investigated alpha and beta diversity of the microbiota from male and female VEH and MOR offspring. There was no significant effect of sex in any of the three alpha diversity metrics so data were collapsed by sex to increase power. No significant effects of age or treatment were noted in Pielou’s evenness (Figure 4A; p=0.1198) or Simpson’s index (Figure 4C; p=0.2744); see Table 1 for complete statistics. However, Shannon’s index (Figure 4B) identified a significant treatment*age interaction (p=0.0178). At P21 and P28, MOR rats had increased Shannon’s index, followed by decreased Shannon’s index at P42 and P56, and no differences at P70. However, given the small magnitude of change and the resolution of any differences by P70, perinatal opioid exposure seems to have only relatively minor effects of alpha diversity.

**Figure 4.**
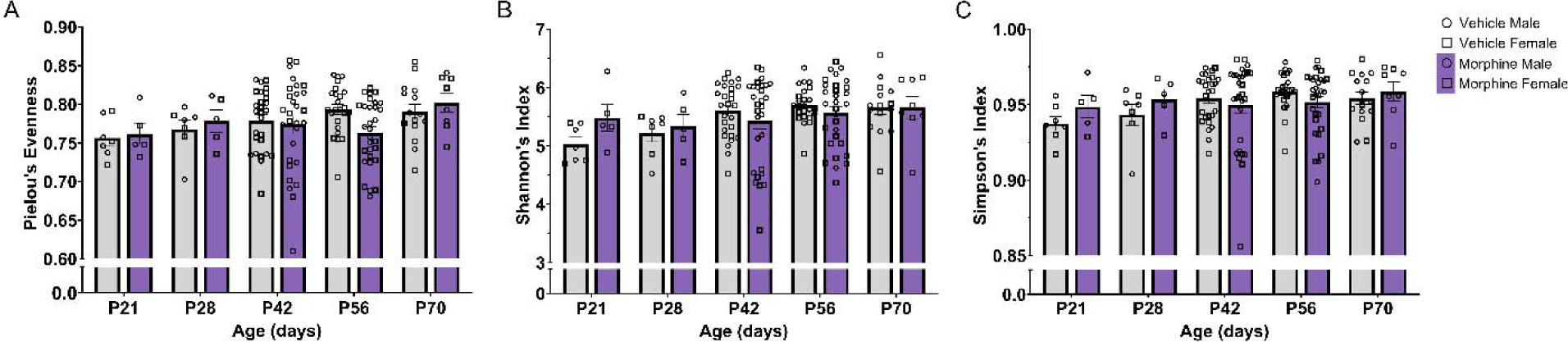
Minor differences were observed in alpha diversity of morphine-exposed offspring. **A.** No significant differences in Pielou’s evenness. **B.** Significant treatment*age interaction in Shannon’s index. C. No significant differences in Simpson’s index. N_VEH_ = 7-27, N_MOR_ = 5-30.

We next analyzed beta diversity separately at P21, P28, P42, P56, and P70 using both Bray-Curtis and unweighted Unifrac (Figure 5). At P21, both Bray-Curtis (p=0.01; Figure A) and unweighted Unifrac (p=0.023; Figure 5B) identified significant differential clustering based on treatment and sex, although no individual comparisons of treatment/sex groups reached statistical significance. At P28, neither Bray-Curtis (p=0.071; figure 5C) nor unweighted Unifrac (p=0.2; Figure 5D) identified significant differences in beta diversity; however, when sex was collapsed, there was a significant effect of treatment for both metrics (p=0.001 and p=0.003, respectively). At P42 (Bray-Curtis, p=0.001, M: q=0.0165, F: q=0.0160; unweighted Unifrac, p=0.001, M: q=0.0015; F: q=0.0015), P56 (Bray-Curtis, p=0.001, M: q=0.0015, F: q=0.0015; unweighted Unifrac, p=0.001, M: q=0.004, F: q=0.003), and P70 (Bray-Curtis, p=0.001, M: q=0.0075, F: q=0.006; unweighted Unifrac, p=0.001, M: q=0.024, F: q=0.03), both Bray-Curtis and unweighted Unifrac identified significant differences in beta diversity for both males and females (Figures 5E-J). In summary, across all five ages sampled, beta diversity was significantly different between VEH and MOR rats, with effects of sex at some ages. Interestingly, although MOR rats were only exposed indirectly to opioids until P7, the observed changes in composition were maintained at least until P70 and even expanded with age.

**Figure 5.**
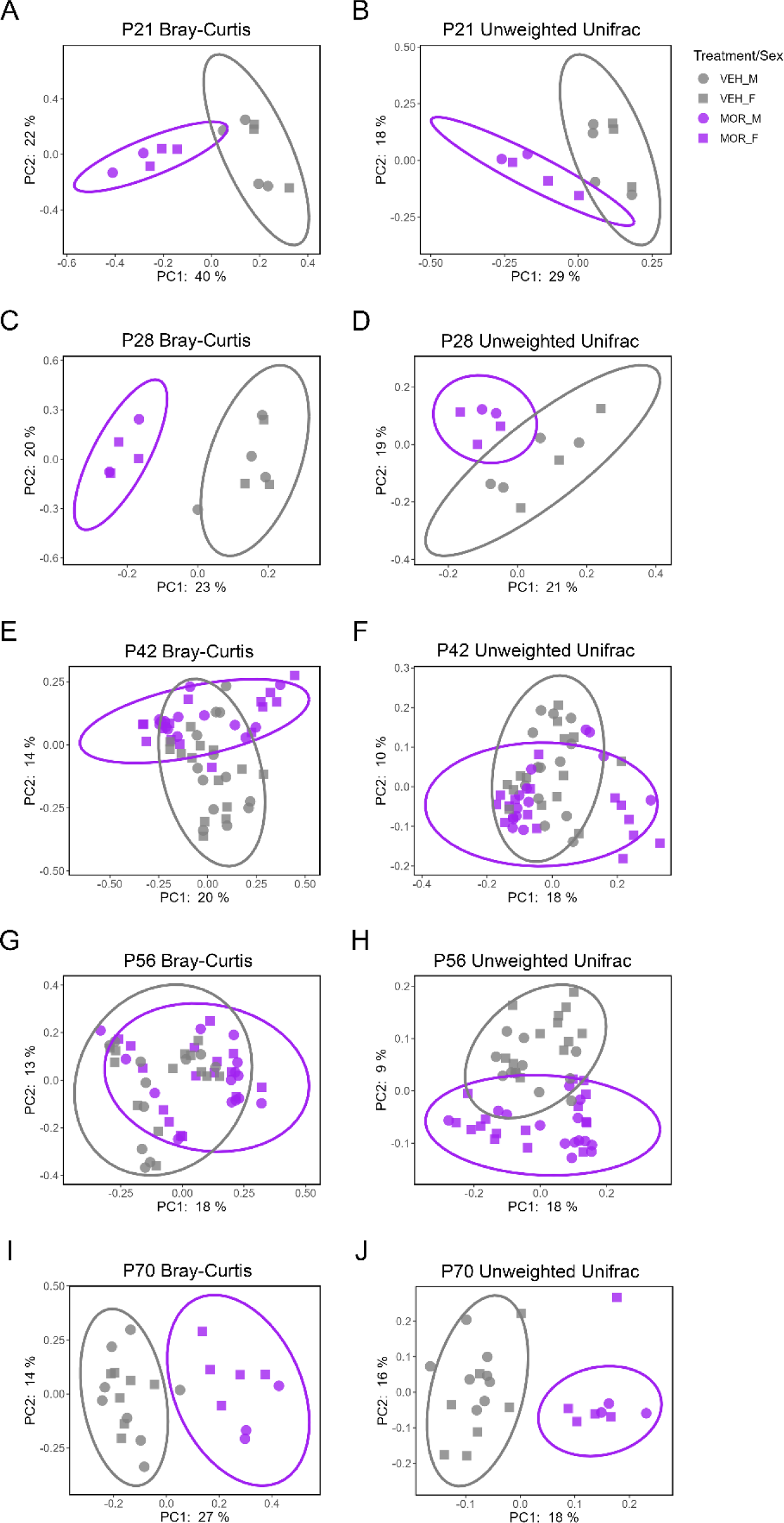
Significant effects of treatment on Bray-Curtis dissimilarity (**A**, P21; **C**, P28; **E**, P42; **G**, P56; **I**, P70) and unweighted Unifrac distances (**B**, P21; **D**, P28; **F**, P42; **H**, P56; **J**, P70).

Hence, we next investigated the relative abundance of specific OTUs responsible for the observed differences in beta diversity noted across age. At P21 (Figure 6A-B), 19 features were differentially abundant (M = 10, F = 6, both = 3), and all 19 were elevated in MOR rats. Of the differentially abundant features, 63.6% were members of the Clostridiales order, particularly the Lachnospiraceae family. Mixed evidence has been reported on the effects of Lachnospiraceae on host health (Vacca et al., 2020), but many Lachnospiraceae members are involved in short-chain fatty acid synthesis, which is important in health and disease. At P28 (Figures 6C-D), 11 features were differentially abundant (M = 5, F = 4, both = 2), with seven upregulated in MOR rats. The families Lachnospiraceae and S24.7 accounted for over half of the differentially abundant features. Although little is known about the role of S24.7, it is a common member of both the rodent and human gut microbiota (Ormerod et al., 2016) and occupies a unique functional niche in the gut, likely playing a role in health and disease. At P42 (Figures 6E-F), ten features were differentially abundant (M = 3, F = 6, both = 1), all downregulated in MOR rats; these included members of the Lachnospiraceae and S24.7 families. Similar results were observed at P56 (Figures 6G-H). Ten features were differentially abundant (M = 6, F = 4), eight of which were downregulated in MOR rats. Many differentially abundant OTUs were members of the Lachnospiraceae and Ruminococceae families. At P70 (Figures 6I-J), 17 features were differentially abundant (M = 11, F = 5, both = 1), 12 of which are upregulated. More than 75% of those differentially abundant OTUs were members of the Clostridiales order, including Lachnospiraceae and Ruminococcaceae.

**Figure 6.**
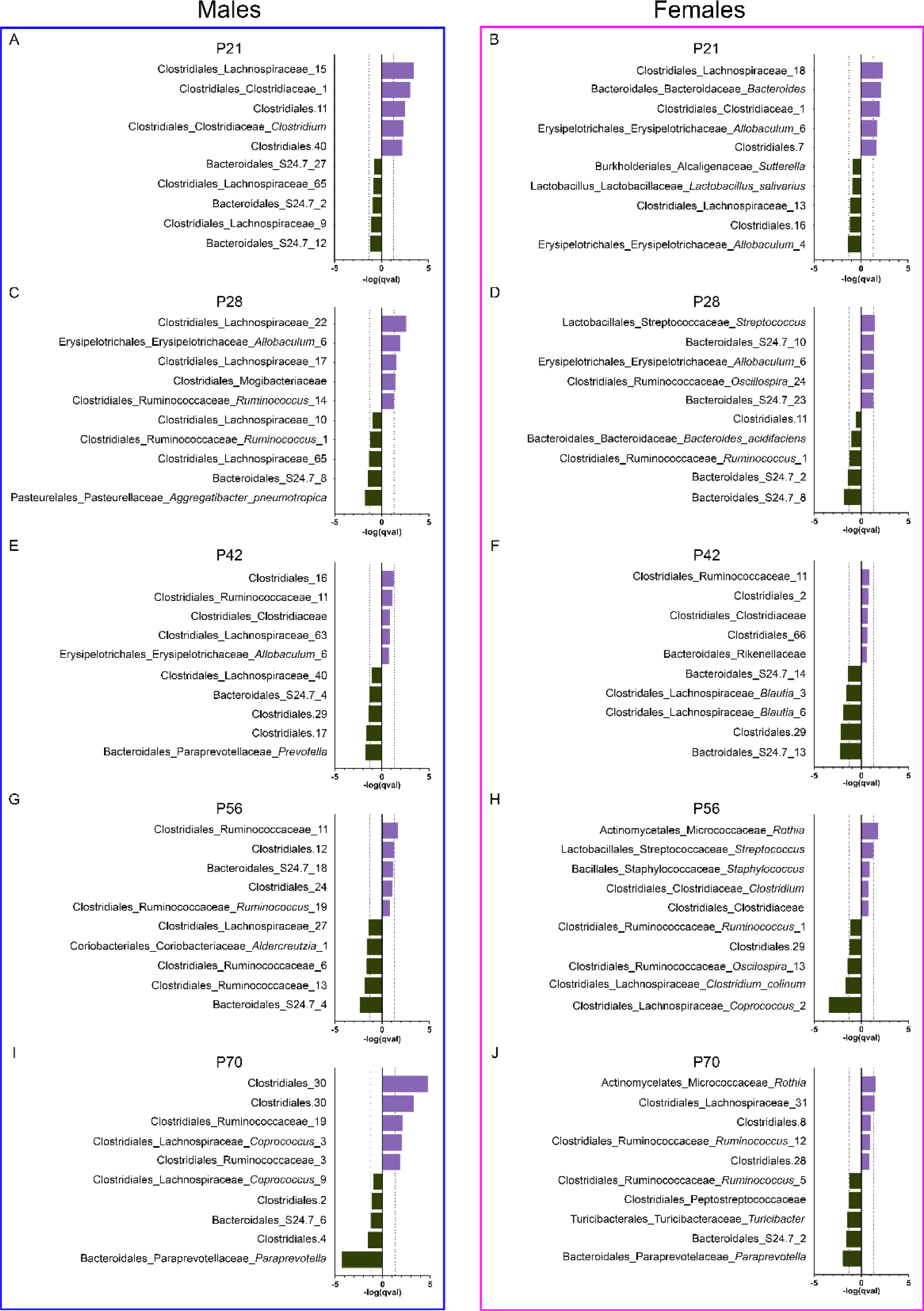
Top five significantly increased and decreased OTUs in the microbiota of morphine-exposed male (**A**, **C**, **E**, **G**, **I**) and female (**B**, **D**, **F**, **H**, **J**) rats at P21 (**A-B**), P28 (**C-D**), P42 (**E-F**), P56 (**G-H**), and P70 (**I-J**). Purple bars represent OTUs increased in the morphine group; gray bars represent OTUs decreased in the morphine group. Dotted vertical lines represent the threshold for significant effect sizes. OTU names = Order_Family_Genera_Species.

Together, this suggests that perinatal exposure to morphine leads to long-term modification of the intestinal microbiota, particularly the Clostridiales order (primarily the Lachnospiraceae and Ruminococcaceae families) and the S24.7 family, both of which were altered at all five timepoints.

### Analysis of microbial maturity

As our analysis included multiple timepoints, we next investigated microbial maturity using the maturity-index function, which trains a regression model on a subset of VEH composition data to compare composition changes as rats age. Maturity index z-scores (MAZ scores) are then calculated to compare the relative maturity between VEH and MOR rats, with negative MAZ scores indicating relatively “immature” intestinal microbiota. MAZ scores were higher for MOR rats at P21 and P28, with average z-scores of 2.74 and 0.71, respectively. However, at P42, P56, and P70, MAZ scores were considerably lower, with average z-scores of −0.69, −1.57, and −1.73, respectively (**Figure 7A**). This suggests that while MOR rats seem to inherit a more mature microbiota, as they age, their microbiota composition does not seem to mature in a comparable manner to VEH rats. We also generated a predicted age based on microbiota composition and compared that to the actual age of the rat at sampling. Positive values indicate a relatively more “mature” intestinal microbiota composition than the actual age at sampling; negative values indicate relatively immature samples. At both P21 and P28, MOR microbiota composition appeared more mature, with an average predicted age of 5.45 and 6.66 days older than the actual sampling age. However, at P42, P56, and P70, MOR microbiota composition was less mature, with an average predicted age difference of 2.86, 6.27, and 9.38 days younger than the actual sampling age (**Figure 7B**). This suggests that as MOR rats reach adulthood, their intestinal microbiota composition remains less mature.

**Figure 7.**
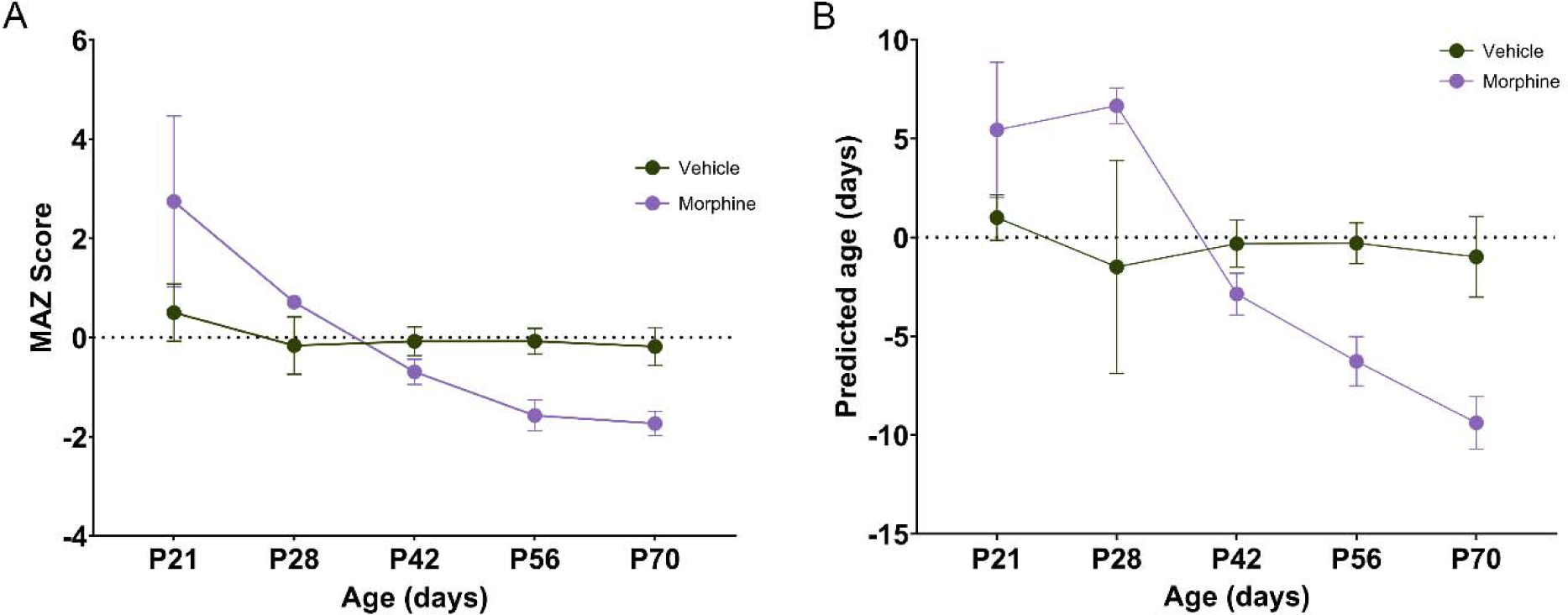
Microbial maturity is altered in morphine-exposed offspring. **A.** MAZ scores for morphine-exposed rats are higher at P21 and P28 but lower at P42, P56, and P70. **B.** Predicted microbial age is higher for morphine-exposed rats at P21 and P28 but lower at P42, P56, and P70. N_VEH_ = 3-15, N_MOR_ = 5-30.

We next applied a linear mixed effects model in order to determine if treatment, sex, or age were significant predictors of maturity index z-scores. As expected, age was a significant predictor of maturity index z-scores (p<0.001) along with treatment (p=0.01). However, sex was not a significant predictor (p=0.608).

## Discussion

The present study was designed to investigate the potential impact of perinatal morphine exposure on gut microbiota composition. We hypothesized that as a result of vertical transmission of a dysbiotic microbiota along with the direct action of opioids on the perinatal gut, the gut microbiota composition of morphine-exposed offspring would be dysbiotic, potentially contributing to the immune dysregulation previously reported in both clinical and preclinical studies of POE (Arter et al., 2021; Harder et al., 2023; Uebel et al., 2015; Witt et al., 2017). Here, we report that morphine administration to dams across pregnancy had a minimal effect on gut microbiota composition; in contrast, long-term and robust changes in gut microbiota composition were observed in male and female MOR offspring.

In our initial studies, we investigated intestinal microbiota composition in morphine- and vehicle-treated dams, as the maternal microbiota has a major influence on the microbiota composition of the offspring. Unexpectedly, no significant differences in alpha diversity were noted in dams treated with morphine across gestation. Rather, alpha diversity generally decreased in both vehicle- and morphine-treated dams as a function of time pregnant, suggesting that pregnancy may have a greater impact on microbiota composition than morphine treatment, at least at the doses tested in this study. Gut microbiota composition was previously found to be dramatically altered across pregnancy: at initiation, composition remains similar to the non-pregnant state, but as pregnancy progresses, the abundance of proinflammatory OTUs increases (Edwards et al., 2017). These physiological changes in the gut microbiota are theorized to alter maternal metabolism in order to support and maintain the pregnancy; however, although not observed here, excessive proinflammatory microbial expansion is associated with a variety of negative outcomes for both the mother and fetus (Edwards et al., 2017). Significant differences were noted in beta diversity in both Bray-Curtis dissimilarity and unweighted Unifrac distance; however, only two OTUs were differentially abundant between vehicle- and morphine-treated dams. Both significant differences were observed before breeding (10 mg/kg), suggesting that morphine’s largest impact on the microbiota is at initiation, and not maintenance.

Despite relatively minor differences observed when comparing vehicle- and morphine-exposed dams, morphine-exposed offspring had significant changes in beta diversity, which were maintained into adulthood and thus, likely permanent. For alpha diversity, although no differences were observed in Pielou’s evenness or Simpson’s index, there were minor (albeit significant) differences in Shannon’s index, although these changes were mainly resolved by P70. Despite only two OTUs showing significant changes in morphine-exposed dams, robust and prolonged differences were observed in the offspring. Together, this suggests that although maternal microbiota composition is one driver of offspring composition, morphine can also act directly on the perinatal gut to alter permeability and peristalsis, creating a conducive environment for microbes to flourish. Interestingly, these changes were maintained into adulthood (P70), despite opioid exposure ending at P7, suggesting that morphine induced long-term damage to the gut and microbiota.

The OTUs differentially expressed in the morphine-exposed offspring over all five ages examined generally belonged to the families Lachnospiraceae, Ruminococcaceae, and S24-7. Although studies have implicated these families in both a health- and disease-promoting role, alterations in such highly abundant members of the microbiota suggest that these changes provide an underlying mechanism whereby infants exposed to opioids *in utero* show a compromised immune response, resulting in higher rates of infection and hospitalization (Arter et al., 2021; Uebel et al., 2015; Witt et al., 2017). Previous studies in adults utilizing chronic opioid exposure report that transplantation of morphine-associated microbiota recapitulates opioid-induced gut pathologies and alterations in immunity (Banerjee et al., 2016). Future studies on perinatal opioid exposure should consider the causal role of gut microbiota composition on opioid-associated immune function by utilizing fecal transplant studies, which may eventually facilitate the development of treatment strategies for children exposed to opioids *in utero*.

Although the mechanism by which opioid-exposed microbiota influences immune system development is unknown, the families Lachnospiraceae and Ruminococcaceae are implicated in bile acid and short chain fatty acid production (Murakami et al., 2018; Vacca et al., 2020), both of which are essential modulators of gut health, microbiota composition, and inflammation. As the majority of OTUs dysregulated in the current study were members of the Lachnospiraceae and Ruminococcaceae families, this suggests that perinatal opioid exposure directly influences bile acid and short chain fatty acid production, paralleling previous work examining chronic opioid exposure in adults (Banerjee et al., 2016; Cruz-Lebrón et al., 2021).

Our results demonstrating a profound and likely permanent impact of perinatal morphine on gut microbiota composition are poised to have a significant impact on the field. Importantly, this study is the first to use a clinically-relevant administration protocol that recapitulates the opioid use profile of pregnant women, including both prenatal and gestational opioids, pulsatile dosing, and increased dosing through pregnancy to account for tolerance. In addition, although previous studies generally report changes in Lachnospiraceae (Grecco et al., 2021), *Lactobacillus*, *Ruminococcus*, *Allobaculum* (Abu et al., 2021), and Clostridia (Lyu et al., 2022), all of which were also altered in the current study, those studies are limited in scope, typically sampling offspring at one age (Grecco et al., 2021; Lyu et al., 2022) or timepoints limited to weaning (Abu et al., 2021) when the gut microbiota composition is still in flux as a result of diet change. Our studies are the first to report that opioid-induced gut microbiota dysbiosis is not only present during early life, but is maintained into adulthood.

In the current study, we also report that morphine-exposed male and female rats had significantly lower microbial maturity in adulthood. Lower microbial maturity is generally defined as a distinct microbial state, rather than a lack of maturity (Subramanian et al., 2014), meaning that MOR rats have distinct composition and developmental trajectory. Lower maturity has been reported in severely malnourished children and was maintained over months despite the introduction of therapeutic food (Subramanian et al., 2014). This suggests that early perturbations of gut microbiota development are maintained long-term. Furthermore, children born premature and admitted to the neonatal intensive care unit had lower microbial maturity vs. healthy controls, and lower microbial maturity was associated with higher beta diversity volatility in early childhood (Yee et al., 2019). Beta diversity volatility, or the change in beta diversity over time, is considered a hallmark sign of disease, as the microbiota should generally remain stable after weaning and solid food introduction (Bokulich et al., 2018). Together, this suggests that morphine-exposed male and female rats have lower microbial maturity, and that this lower maturity may be associated with an increased disease risk and beta diversity volatility.

The results of this study have significant clinical implications for children born with NOWS, who often suffer from gastrointestinal complications and are at higher risk for infection later in life. Though it is currently theorized that *in utero* opioid exposure alters gut microbiota composition in humans (Maguire and Gröer, 2016), to date, no clinical studies have compared the microbiota of children exposed to opioids *in utero* to healthy controls. As a consequence, the mechanism by which opioid-exposed microbiota influences human health or increases disease risk later in life is currently unknown.

In summary, this study provides evidence that perinatal morphine leads to long-term modification of the intestinal microbiota, in both male and female offspring, with a potential impact on the immune system later in life (Harder et al., 2023). Specifically, we report here that members of the families Lachnospiraceae, Ruminococcaceae, and S24-7 are altered until at least P70 and that the maturity of the morphine-exposed microbiota composition is lower than that of vehicle rats in adulthood. The use of a clinically translatable model of *in utero* opioid exposure provides initial evidence to support the hypothesis that human infants born with NOWS suffer from gut dysbiosis, potentially leading to long-term negative health outcomes and increased risk for infection and hospitalization.

## Acknowledgments

This work was supported by the National Institutes of Health (1RO1DA041529). BC’s laboratory is supported by a Starting Grant from the European Research Council (ERC) under the European Union’s Horizon 2020 research and innovation program (grant agreement No. ERC-2018-StG-804135). The authors thank the Genom’IC platforms (INSERM U1016, Paris, France) for their help with sequencing approach.

The funding source was not involved in study design, data collection, analysis or interpretation, manuscript writing, or the decision to publish this article. Morphine sulfate was provided by the NIDA Drug Supply Program.

